# TAFKAP: An improved method for probabilistic decoding of cortical activity

**DOI:** 10.1101/2021.03.04.433946

**Authors:** R.S. van Bergen, J.F.M. Jehee

## Abstract

Cortical activity can be difficult to interpret. Neural responses to the same stimulus vary between presentations, due to random noise and other sources of variability. This unreliable relationship to external stimuli renders any pattern of activity open to a multitude of plausible interpretations. We have previously shown that this uncertainty in cortical stimulus representations can be characterized using a probabilistic decoding algorithm, which inverts a generative model of stimulus-evoked cortical responses. Here, we improve upon this method in two important ways, which both target the precision with which the generative model can be estimated from limited, noisy training data. We show that these improvements lead to considerably better estimation of the presented stimulus and its associated uncertainty. Estimates of the presented stimulus are recovered with an accuracy that exceeds that of standard decoding methods (SVMs), and in some cases even approaches the behavioral accuracy of human observers. Moreover, the uncertainty in the decoded probability distributions better characterizes the precision of cortical stimulus information from trial to trial.

## Introduction

Brains and brain researchers face a common challenge: interpreting the information conveyed by neural responses about external stimuli in the world. This is a hard challenge, because stimuli and responses typically do not have a straightforward and unambiguous relationship. Sources of variability and noise, both external and internal to the brain, make it so that the exact same external stimulus can lead to a different response each time it is presented to the observer. By the same token, because of neural variability, any given neural response typically has a range of possible causes. Thus, interpreting a neural response is not as simple as applying a function or lookup table to find the single external cause that uniquely explains it. There may be one hypothesis that *best* explains the neural data, but others are usually also plausible. The interpretation, then, is *uncertain*.

This uncertainty is particularly relevant when integrating across different sources of information. For example, when downstream areas integrate information from multisensory sources. When integrating information, it is statistically optimal to take sensory uncertainty into account, giving more weight to more reliable (less uncertain) information in the integration process. Interestingly, human behavior often matches or approximates such statistically ideal behavior^1–3^, suggesting that the brain represents and computes with uncertainty. Precisely how such an ideal inference process is implemented in the brain, however, is currently unknown, and the topic of much recent study^4–7^.

To address the neural code for uncertainty, we recently developed a probabilistic decoding approach (**Fig. 1A-B**) and showed that it successfully characterizes the degree of stimulus uncertainty contained in population activity obtained from human visual cortex^5,8^. The technique tracks the degree of uncertainty associated with cortical stimulus representations on a trial-by-trial basis, and can be used to test hypotheses regarding optimal perceptual inference and decision-making in human observers^5,6^. We will retroactively refer to this decoding method as PRINCE (**Pr**obabilistic **In**ference from activity in **C**ort**e**x).

**Figure 1:**
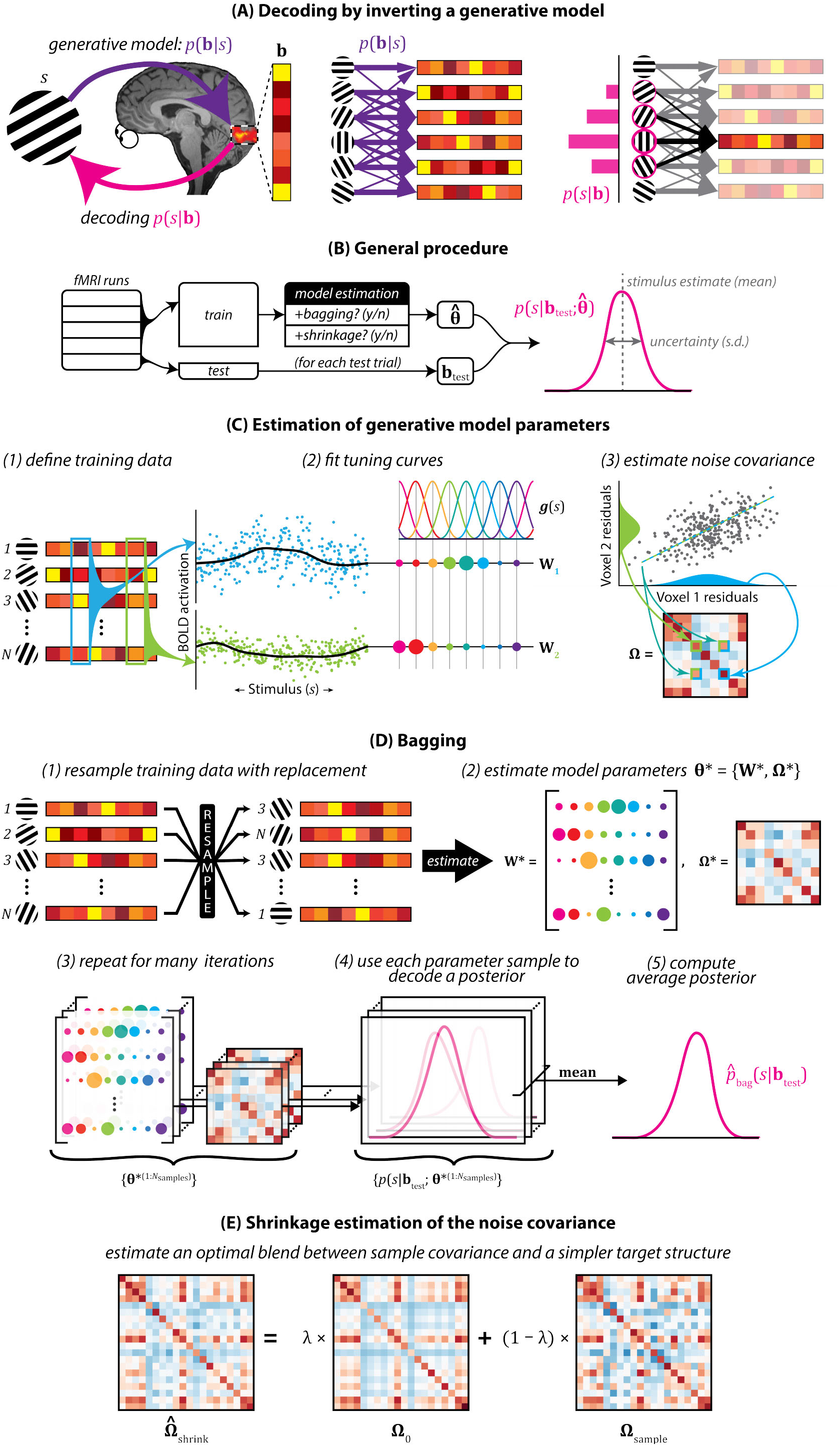
Illustration of the generative model-based probabilistic decoding procedure(s). (**A**) Simplified illustration of the rationale behind the approach. The generative model *p*(**b**|*s*) describes the probability of different patterns of cortical activity for a given stimulus. When decoding a probability distribution over stimuli *p*(*s*|**b**) from a given cortical activity pattern, this generative model is inverted, reasoning backwards from activity to stimuli. The right side of the panel provides a simplified illustration of this principle. For this toy example, the generative model consists of a lookup table of conditional probabilities (purple arrows – thickness indicates probability) between a small set of stimuli and a small set of activity patterns. When one of these patterns is observed, as shown on the far right (observed pattern is shown as opaque), decoding consists of consulting the lookup table to determine for each stimulus the likelihood that it evoked the observed pattern, and then normalizing these probabilities so they form a proper distribution over stimuli (instead of over activity patterns). In reality, the generative model consists of a Normal probability density function (with free parameters fit to training data) to ensure continuous coverage of a much larger space. However, the basic principle is the same. (**B**) General procedure used for all probabilistic decoding approaches discussed in this study. First, fMRI data are split into train and test sets. Train data are used to estimate generative model parameters, with or without bagging, and with or without shrinkage estimation of the covariance matrix. Each activity pattern in the test set is used to decode a posterior distribution given the estimated model parameters. The mean of the posterior serves as the decoded estimate of the presented stimulus. The width (circular standard deviation) of the posterior serves as measure of the degree of uncertainty in the cortical response. (**C**) Estimation of generative model parameters. fMRI data are first divided into train and test sets. Model parameters are estimated from the train data. Using these parameters, posteriors are decoded from activity patterns in the test set. The generative model is specified by the tuning functions and noise covariances of the fMRI voxels. Orientation tuning functions are estimated first. For each voxel, a tuning function is estimated from its responses in the training data, through least-squares regression, as a weighted sum of eight bell-shaped basis functions^22^. This procedure is illustrated for two hypothetical voxels, with filled circles indicating the fitted weights for each of the basis functions. The residual responses around the fitted tuning functions are used to estimate a noise covariance matrix. The diagonal of this matrix contains each voxel’s individual noise variance. The off-diagonal elements encode the noise covariance between each pair of voxels. (**D**) Bootstrap aggregating (bagging) procedure to account for imprecision in the estimated generative model parameters due to noise in the training data. First, training data are resampled by randomly drawing trials (stimuli and activity patterns) with replacement. Resampled training data are then used to estimate a set of model parameters. This is repeated many times. Each so-obtained parameter value is used to decode a posterior from every activity pattern in the test set. For each test trial, this results in a large set of distributions, which are averaged to obtain a single decoded posterior per trial. (**E**) Shrinkage-based covariance estimation flexibly interpolates between a rich but error-prone estimate of the voxel noise covariance (the sample covariance matrix **Ω**_sample_) and a simplistic but more stable estimate (the *shrinkage target* **Ω**_0_). *λ* controls the mixture proportions of these two extremes.

In the current work, we present several substantial modifications to this original approach, and show that the modified algorithm provides significantly better estimates of the degree of uncertainty associated with stimulus representations in cortex. These modifications target the uncertainty in model parameters that arises due to limited and noisy training data. We account for this imprecision in model parameters in two different ways. First, we aggregate information over many possible settings of the generative model parameters, using a sampling-based technique called *bagging*^9^. Second, we more flexibly estimate the spatial noise covariance in cortical activity, using a regularization technique known as “shrinkage”^10,11^. This gives rise to a new version of our decoding approach, called TAFKAP (**T**he **A**lgorithm **F**ormerly **K**nown **A**s **P**rince). Both changes to the algorithm are first validated on synthetic data, and then tested on neuroimaging data from human participants. We find that TAFKAP achieves substantially better accuracy in estimating stimulus orientations and trial-by-trial uncertainty. Of particular interest is the ability to predict human behavior from decoded uncertainty estimates, which also increased markedly, suggesting that our modifications enhance the algorithm’s sensitivity to the degree of uncertainty contained in neural populations underlying the fMRI voxels.

## Methods

### Participants

All subjects provided written and informed consent prior to participation. Participants were healthy, adult volunteers (seven female, eleven male), aged 22-31 years, with normal or corrected-to-normal vision. The study was approved by the Radboud University Institutional Review Board.

### MRI data acquisition

fMRI data were originally collected in the context of a different study^5^. They were acquired using a Siemens 3T Magnetom Trio scanner located at the Donders Center for Cognitive Neuroimaging, using an eight-channel occipital receiver coil. Functional images (T2*-weighted gradient-echo EPI sequence; TR: 2000 ms; TE: 30 ms; flip angle: 90°; 2.2 mm isotropic voxels; FOV: 64 × 64) consisted of 30 slices, aligned perpendicularly to the calcarine sulcus, and covered all of occipital and some of parietal and temporal cortex. A high-resolution T1-weighted anatomical scan (magnetization-prepared rapid gradient-echo (MPRAGE) sequence; 1 mm isotropic voxels; FOV 256 × 256) was also acquired at the start of each scan session.

### Experimental design & stimuli

Participants completed between 10-18 runs of an orientation estimation task inside the MRI scanner. Runs consisted of 18 trials. Each trial started with the presentation of a sinusoidal grating stimulus (duration: 1500 ms; spatial frequency: 1 cycle/degree; random spatial phase; 2 Hz sinusoidal contrast modulation; peak contrast: 10%) presented inside an annular window centered on fixation. The annulus had an inner radius of 1.5 degrees, an outer radius of 7.5 degrees, and grating contrast decreased linearly to 0 over the outer and inner 0.5 degrees of its radius. The orientation of the stimulus was determined (pseudo-)randomly to ensure that orientation space was sampled roughly evenly in each run. The stimulus was followed by a retention interval (6500 ms) after which participants were prompted to report the previously seen orientation, by rotating a bar that appeared at the center of the screen (width: 0.1 degrees; length: 2.8 degrees). Participants adjusted the orientation of the bar via an MRI-compatible button box, pressing separate buttons for clockwise or counterclockwise rotation. The bar remained on screen for a total of 4 seconds, fading into the background during the last second of this window to alert the participant that the response window was ending. After a 4-s interval, the next trial was initiated by the presentation of another orientation stimulus. Throughout the run, participants were instructed to maintain fixation on a centrally presented, black-and-white bullseye target (radius: 0.25 degrees).

Each participant also completed two runs of a visual localizer task, in which 100%-contrast flickering checkerboard stimuli (check size: 0.5 degrees) were presented within the same annular window as the orientation stimuli. Stimuli were presented in blocks of 12 seconds, alternating with fixation blocks of equal length. During stimulus blocks, a new, random checkerboard pattern was presented every 100 ms. In addition, a retinotopic map of each participant’s visual cortex was acquired in a separate scan session, using standard retinotopic mapping procedures^12–14^.

Participants viewed visual stimuli inside the MRI scanner through a mirror mounted on the head-coil. Stimuli were generated using a Macbook Pro computer running MATLAB and the Psychophysics Toolbox^15,16^, and displayed on a rear-projection screen using a luminance-calibrated projector (EIKI; resolution 1024 × 768; 60 Hz refresh rate)

### Behavioral data

Behavioral errors were computed as the acute-angle difference between the presented and reported orientations on a given trial. With an average error of 6.29 ± 0.25° (mean ± SEM across participants), subjects generally performed well on this task. Before further analysis, behavioral errors were corrected for an orientation-dependent bias. Each observer’s behavioral errors were fit separately with a 4-degree polynomial curve as a function of stimulus orientation, and the residuals from this fit were used in subsequent analyses (e.g. when computing behavioral variability). Trials where the corrected behavioral error exceeded three standard deviations from an observer’s mean error were considered random guesses and excluded from further analysis – this resulted in a maximum of 6 trials being excluded for one observer, and between 0-3 trials for the remaining observers.

### fMRI pre-processing and ROI selection

Regions of interest (ROIS; V1, V2 and V3) were defined on the reconstructed cortical surface using standard procedures^12–14^. Within each of these areas, we selected for subsequent analysis all voxels that responded significantly to an independent visual localizer stimulus, using a lenient statistical threshold (*p* < 0.05, uncorrected). Functional data were analyzed in the native space for each participant.

Functional images were motion-corrected using FSL’s MCFLIRT algorithm^17^, and aligned to a previously collected anatomical reference scan using FreeSurfer^18^. BOLD time series were high-pass filtered, with a cut-off period of 40 s, to remove slow temporal drifts from the signal. No slice timing correction was performed. Residual motion-induced fluctuations in the BOLD signal were removed using linear regression (using 18 motion regressors, consisting of the 6 rotation and translation estimates generated by the MCFLIRT algorithm for each time point, as well as the squares and temporal derivatives of these values). This nuisance regression also included a predictor constructed from the mean signal intensity across all voxels that fell within retinotopically labeled cortical regions, which removed global fluctuations from the BOLD response.

Voxel timecourses were Z-normalized separately for each fMRI run. Specifically, the activity of the *i*-th voxel in the *n*-th time point within each trial was normalized by the mean and standard deviation of that voxel’s responses in the *n*-th time points of all trials in the same run. To obtain a single response measure for each voxel on each trial, these Z-scored values were shifted by 4 s of hemodynamic lag, and then averaged across the first 4 s of each trial. This time window ended 8 s after stimulus onset (i.e. at the start of the response window) to ensure that no activity from the response window could influence the decoding analyses.

### Probabilistic decoding procedures

The new probabilistic decoding algorithm TAFKAP includes two main modifications with respect to its predecessor^5^: bootstrap-aggregation (*bagging*) across the training data, and regularized estimation of the voxel noise covariance matrix. Each of these modifications was validated separately on synthetic data. Thus, a total of four different probabilistic decoding procedures are considered here: (1) PRINCE, (2) PRINCE+bagging, (3) PRINCE+shrinkage and (4) TAFKAP – the full new version of the algorithm with both modifications.

All four of these procedures followed the same general approach (**Fig. 1**). A generative model is first estimated using some training data, and then this estimated model is inverted to decode a probability distribution from test data that was withheld during training. This final decoding step is basically identical between all decoders, which differ mainly in the way that the generative model is estimated. We first explain the generative model and overall cross-validated decoding procedure, and then describe how the parameters of the generative model are estimated for each decoder.

#### Generative model

The approach starts with the assumption that cortical activity varies randomly from trial to trial around a fixed orientation tuning curve that is different for each fMRI voxel^5,8^:

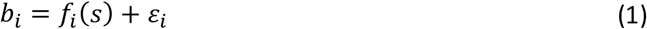

where *b*_*i*_ is the response of the *i*-th voxel, which is a sum of its tuning curve *f*_*i*_(*s*) to stimulus orientation *s* (in °) and random noise *ε*_*i*_. Orientation tuning curves are modeled as a linear combination of 8 bell-shaped basis functions:

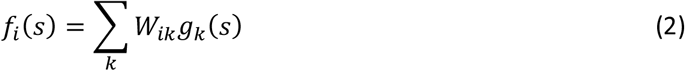

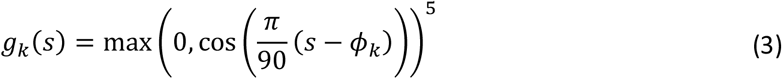

where *g*_*k*_(*s*) is the *k*-th basis function, which peaks at orientation *ϕ*_*k*_, and *W*_*ik*_ scales that basis function’s contribution to the *i*-th voxel’s tuning curve. Peak orientations *ϕ*_*k*_ are regularly spaced in the interval [0,180]°.

Around its tuning function *f*_*i*_(*s*), any given voxel’s response is assumed to vary randomly due to Normally distributed noise **ε**. This noise is assumed to be correlated between voxels, with covariance **Ω**, such that 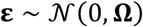. The probability of a cortical activity pattern **b** = [*b_i_*]^T^ is therefore given by:

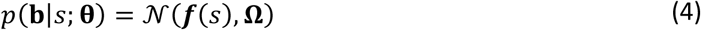

where ***f***(*s*) = [*f*_*i*_(*s*)] are the values of the voxel tuning functions, and **θ** describes the free parameters in the model. Specifically, **θ** = {**W**, **Ω**}, where voxel weights **W** and covariance structure **Ω** are determined by the data.

#### Cross-validated training & testing

Model parameters (**θ**) were estimated from the data in a leave-one-run-out cross-validation procedure. That is, for each participant, fMRI datasets were divided into training and testing partitions (**Fig. 1B**), such that each fMRI run served as the test set exactly once. Training data were used to estimate model parameters, while data from the test set was used to decode posterior distributions from the activity patterns on a trial-by-trial basis. The way in which model parameters were estimated depended on the specifics of each decoding procedure.

After obtaining estimates of all free parameters in the generative model, posteriors are decoded from activity patterns in the test set by applying Bayes’ rule:

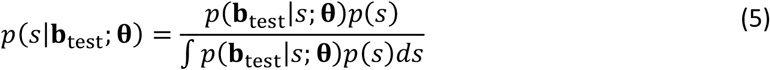

where we use a flat prior *p*(*s*) (reflecting the uniform distribution of stimuli in the experiment), and the normalization constant in the denominator is computed numerically. We use a discrete approximation for the posterior, evaluating equation (5) at 100 equally spaced orientations. The circular mean of the posterior serves as our estimate of the presented stimulus, and its circular standard deviation as a measure of the degree of uncertainty in this estimate (**Fig. 1B**).

#### Model estimation: PRINCE

In PRINCE, our original decoding procedure^5,8^, model parameters are estimated in two steps (**Fig. 1D**). First, orientation tuning curves are estimated through an ordinary least-squares regression:

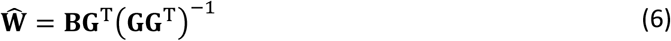

where 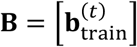 is a matrix of all activity patterns in the train set (where *t* indexes trials), and 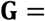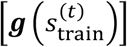 are the values of the basis functions for the stimuli presented on the same trials. In addition, we also included in **G** a set of nuisance regressors, designed to capture any residual activation to the stimulus presented on the preceding trial. These regressors are constructed by shifting each column 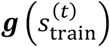 by a single trial (for trials that were the first in an fMRI run, there was no immediately preceding stimulus, and so these values were set to 0). Note that decoded posteriors were based only on the current trial-tuning curves, and not on the part of the response that could be explained by the preceding stimulus.

Next, we estimate the voxel-by-voxel noise covariance matrix. This covariance is assumed to have a simple structure:

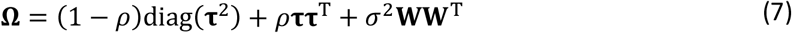

where 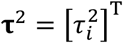 are variance parameters of noise that is independent between voxels, *ρ* is a global correlation parameter for noise that is shared among all voxels, and *σ*^2^ is a variance parameter for noise that is shared between voxels with similar orientation tuning properties (**WW**^T^ can be thought of as a matrix representing the degree of similarity in tuning). The parameters of this covariance model (**τ**, *ρ* and *σ*) are found by numerically maximizing their likelihood under the training data, conditioned on the estimate of **W**.

#### Model estimation: TAFKAP

The new TAFKAP decoding procedure is mostly identical to that used in PRINCE, except that it alters the estimation of the generative model in two ways, adding **S**ampling and **S**hrinkage techniques to better account for the uncertainty that automatically arises when estimating a generative model from limited and noisy training data.

#### TAFKAP: bagging

TAFKAP employs a sampling method to obtain model parameters, which is called bootstrap-aggregating, or *bagging*. It is implemented by randomly resampling training data with replacement many times, and estimating generative model parameters from each resampled dataset (**Fig. 1D**). Specifically, for each bootstrap sample *j*, a random list (of length *N*_traintrials_) of training trial indices is generated, which may include repetitions (and omissions). A resampled set of training data {**B**^∗(*j*)^, **s**^∗(*j*)^} is then created, by selecting the rows of **B** (voxel responses) and **s** (stimuli) corresponding to these trials. These resampled training data thus have the same dimensions as the original training data. Model parameters **θ**^∗(*j*)^ = {**W**^∗(*j*)^, **Ω**^∗(*j*)^} are then estimated from the resampled dataset, either using the original estimation procedure (see previous section) or a modified version with covariance shrinkage (described in the next section).

For each obtained parameter sample **θ**^∗(*j*)^ and test pattern, we then compute a posterior *p*(*s*|**b**_*test*_, **θ**^∗(*j*)^). Across bootstrap samples, posteriors are subsequently averaged to obtain one decoded posterior for each test trial:

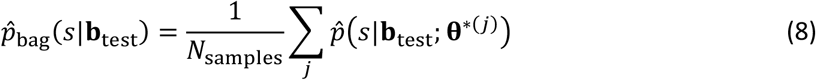

*N*_samples_ can either be fixed to a pre-set value, or it can be determined while sampling, based on a convergence criterion. In the simulation analysis, we used a fixed setting of 10,000 samples. When decoding real fMRI data, this same value was used as a maximum, but the algorithm was also allowed to terminate earlier, if the running estimates of the test-trial posteriors had not changed substantially in the last epoch of 100 samples (specifically, if the largest Jensen-Shannon divergence between the current and previous estimates of these posteriors was less than 10^−8^).

This bagging procedure enables one to average over many possible model parameters – all of which are plausible under the training data, rather than having to commit to a single parameter value. In addition to these parameter values, we also applied sampling techniques to the placement of the orientation basis functions (while the shape of the basis functions themselves remained constant across samples). That is, in the original decoding procedure, the basis functions are placed such that the first basis function peaks at 0°, and the others follow at equal intervals of 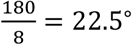. This is a reasonable choice, but other settings are also sensible. For instance, one could shift the entire set of basis functions by half their spacing, such that the first function is centered on an orientation of 11.25° (while keeping functions spaced at 22.5°). Thus, the exact placement of the basis functions is somewhat arbitrary, and the optimal placement for estimating voxel tuning functions might not be constant across voxels. The bagging procedure used in TAFKAP therefore implements the option to switch randomly between four different basis sets: the original basis (with the first function centered at 0°) and three shifted copies (with the first function centered at 5.625°, 11.25° or 16.875°, respectively). At the start of each bootstrap iteration, one of these sets is randomly selected, before proceeding with the estimation of the model parameters. In the current study, this basis-shifting feature was included in the decoding analysis of real fMRI data (i.e. by alternating between four equally spaced offsets), but not for the analysis of simulated data (for which the true (simulated) basis was known by construction).

#### TAFKAP: covariance shrinkage

The second of the two changes applied to the model estimation step is regularized estimation of the voxel noise covariance matrix. This procedure blends the simplified covariance structure used in PRINCE (equation (7)) with the sample covariance matrix of the training data (**Fig. 1E**). That is, the simpler structure now acts as a target to which the sample covariance is “shrunk”^11,19^:

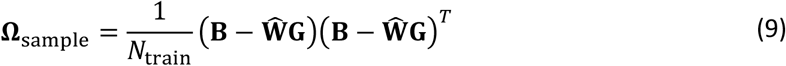

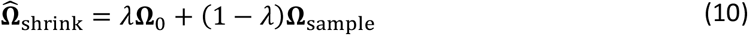

where *λ* controls the overall degree of shrinkage, between [0,1], and *N*_train_ is the number of trials in the training set from which the covariance is estimated. The shrinkage target was identical to the covariance structure of the original model, except for two small modifications:

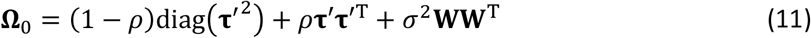

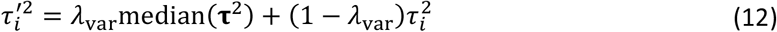

That is, the diagonal of variances in **Ω**_0_ is itself shrunk towards its median value, by an amount controlled by *λ*_var_ – an approach modeled after [11]. Another difference is in the way the parameters of **Ω**_0_ are estimated. In PRINCE, we used an iterative optimization procedure to find the maximum-likelihood estimates of these parameters. This optimization procedure can take a long time to converge (on the order of hours or days, depending on the number of voxels in the data set), which is feasible when parameters are estimated only once for any given set of training data. When using the bagging approach described previously, however, parameters are estimated thousands of times: once for each bootstrap resampling of the training data. In this case, a lengthy iterative procedure for each of these estimations is not practically feasible. We therefore abandoned the iterative procedure in favor of a much faster OLS regression approach, which computes the estimate 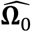 that has the smallest squared deviation to the sample covariance matrix:

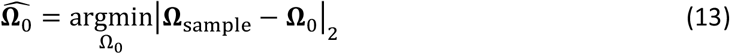

The optimal values of the shrinkage hyperparameters *λ* and *λ*_var_ are selected using a standard, additional leave-one-run-out cross-validation procedure within the training data, also used in [11]. Briefly, the goal is to find those shrinkage values that are likely to provide the best covariance matrix estimate when applied to the test data. This is achieved by trying many settings of shrinkage strengths, and estimating their generalization performance across different splits of the training data. The shrinkage setting with the best performance on the train data is then used to compute the value of 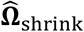 for decoding the test data.

A separate hyperparameter search is performed for each set of training data. For decoding procedures that include bagging, the hyperparameter search is performed on the original (non-bootstrap-resampled) training data. The shrinkage values found in this search are then used for covariance estimation on each bootstrap sample drawn from this training data.

### SVM decoder

To compare our decoding approach to other methods that are widely used in the field of (non-probabilistic) fMRI decoding, we also implemented a decoding procedure based on support vector machines (SVMs). These are most commonly used for discrete classification, but a version called *support vector regression*^20^ (SVR) adapts the SVM method to work for continuous estimation problems like ours. Here, we used the `fitrlinear` function from Matlab’s Statistics and Machine Learning Toolbox, with the ‘Learner’ parameter set to ‘svm’. To accommodate the fact that orientation space is circular, we fit two separate linear SVR models, to predict the sine and cosine of the presented orientation:

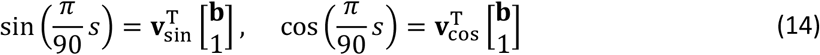

where **v**_sin_ and **v**_cos_ are weight vectors (of size (*N*_voxels_ + 1) × 1, with the last entry modeling an intercept term), and the multiplication by 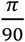 transforms the stimulus orientation in degrees to the circular interval [0, 2*π*]. These weight vectors are fit to a set of training data using the SVR learning algorithm. Given an activity pattern from a test set, these vectors are then used to predict the sine and cosine of the orientation presented on that trial:

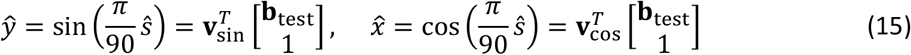

Finally, these predicted values are transformed back to the orientation domain:

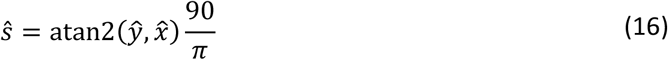

where atan2(∙) is the two-argument arctangent function. This estimate 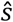 serves as the decoded orientation for that trial. Note that this SVM-based decoding approach gives a single orientation (point) estimate per trial, and does not produce estimates of uncertainty. To create independent train and test sets, we used the same leave-one-run-out cross-validation procedure as for the other decoding methods, such that each fMRI run served as the test set once.

### Simulations

The bagging and covariance estimation techniques described above were first tested separately on simulated data. Eighteen synthetic fMRI datasets were simulated based on distinct randomization seeds. BOLD responses were simulated for a realistic number of train trials (250 or 270) and a large number of test trials (500), for a population of 50 voxels. To ensure a realistic distribution of signal and noise, each dataset was simulated based on estimated voxel tuning curves and a noise covariance matrix from one of the human participants in our study (such that each simulated dataset was based on a different participant). Specifically, for each participant, we selected the 50 voxels from V1, V2 and V3 that were most strongly activated by the independent localizer stimulus. From these voxels’ responses across all fMRI runs, we estimated tuning coefficients using an OLS regression (equation (6)), and computed the sample covariance of the regression residuals (equation (9)). We denote the obtained values **W**^(sim)^ and **Ω**^(sim)^, respectively. These parameter values were then used to generate data for a simulated participant. Stimulus orientations for the train and test trials were drawn randomly from a Uniform distribution, and voxel responses were simulated as random draws from the multivariate Normal distribution specified in equation (4) with mean defined by **W**^(sim)^, and covariance **Ω**^(sim)^.

### Statistical analyses

#### Simulations

We first compared the four decoding procedures on simulated data (i.e., PRINCE, PRINCE+bagging, PRINCE+shrinkage & TAFKAP), by evaluating the quality of their decoded posteriors. For simulated data, decoded posteriors can be compared to the true posterior encoded in a pattern of voxel responses; this posterior is determined by the generative model that was used to simulate these data. Specifically, the true posterior is computed by evaluating equation (5), with **θ** = **θ**^(sim)^ = {**W**^(sim)^, **Ω**^(sim)^}. We denote this posterior by *p*_true_(*s*|**b**_test_). Each decoded posterior’s fidelity was quantified by computing its Kullback-Leibler divergence to the true posterior:

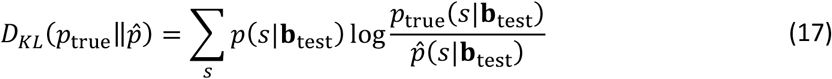

where 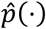 denotes a posterior estimated with one of the four decoding methods. KL-divergences between estimated and true posteriors were statistically compared using paired *t*-tests.

#### fMRI data

In the analyses of real fMRI data, we compared our original decoding procedure (PRINCE) to the new version (TAFKAP), with both modifications (bagging and shrinkage) included. To evaluate the extent to which each decoding approach accurately captured the information contained in fMRI activity patterns from human observers, we assessed (1) how well the decoded posterior predicted the stimulus presented on that trial, and (2) how well the decoded posterior quantified the precision of information in the cortical stimulus representation. For the first metric, we compared the true stimulus orientation to the stimulus estimate (posterior mean) from each decoded posterior, computing both the mean absolute error and circular correlation between true and decoded stimulus orientations. For the second, we compared the width of the decoded posteriors to the precision of the decoded stimulus estimates and the behavioral stimulus reports. These analyses are explained in more detail below.

The decoding errors (i.e. the angle between presented and decoded orientation) achieved by the different decoding procedures were compared pairwise within individual subjects, and across subjects. Within each subject, a two-tailed Wilcoxon signed-rank test was performed on the trial-by-trial (absolute) errors from different decoders. Across subjects, a two-tailed Wilcoxon signed-rank tests was performed on the mean absolute error achieved by different decoders.

The decoding correlations (i.e. the circular correlation between presented and decoded orientations) achieved by the different decoding procedures were also compared pairwise within and across subjects. Within each subject, a permutation test was used to statistically assess the difference in correlation between decoders (we used 10,000 random permutations of the decoded orientation values to simulate the null hypothesis that decoded orientations are uncorrelated to the true stimulus orientations). Across participants, two-tailed Wilcoxon signed-rank tests were computed on the correlation achieved by each decoder for each subject.

To examine whether the decoder’s uncertainty estimates reflect the actual uncertainty in cortical activity, we tested the trial-by-trial relationship between decoded uncertainty and variability in decoded orientation estimates, and between decoded uncertainty and variability in the observers’ behavioral orientation reports. We examined this relationship in two ways. First, we used a regression analysis. Specifically, for each participant and decoding approach, trials were divided into four equally-populated bins of increasing uncertainty. Within each bin, we computed the mean decoded uncertainty and the (circular) standard deviation of the (behavioral or decoding) error in orientation estimates. For each decoding analysis, we then fit a multiple linear regression model to these bin statistics, with decoded uncertainty as the regressor of interest. We also modeled a separate intercept regressor for each subject. This was done to ensure that the estimated relationship between uncertainty and error variance reflected the (within-subject) trial-to-trial correlation, rather than a relationship between uncertainty and error across subjects. For plotting purposes (**Fig. 4–5**), these intercepts were removed from the data and replaced by the corresponding group averages. Regression coefficients were transformed to partial correlation coefficients, and their *p*-values were computed by means of a two-tailed *t*-test. As a second approach, we analyzed the relationship between decoded uncertainty and across-trial variability in (behavioral or decoded) orientation estimates by means of a bin-free model comparison. Here, errors were modeled with a Normal distribution, the standard deviation of which depended on decoded uncertainty:

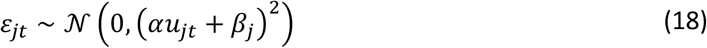

where *j* indexes participants and *t* indexes trials, *ε*_*jt*_ is the decoder or behavioral error, *u*_*jt*_ is the decoded uncertainty, *α* is the slope of the relationship between decoded uncertainty and error standard deviation, and *β*_*j*_ is the intercept of subject *j*’s error variance. This model formalizes the hypothesis that there is a trial-by-trial correlation between decoded uncertainty and the width of the across-trial error distribution. The model was fit to the data (errors and uncertainties) by numerically maximizing the likelihood of its parameters (*α* and {*β*_*j*_}), with *α* constrained to be strictly positive. The goodness-of-fit of the model was quantified by means of the Bayesian Information Criterion (BIC)^21^. To determine whether decoded uncertainty was reliably linked to error variability, this BIC was compared to that of a null-model, in which no such relationship between decoded uncertainty and error variability existed (equivalent to setting *α* = 0 in equation (18)). To adjudicate between the PRINCE and TAFKAP decoding algorithms, we compared BIC values for models fit with uncertainty values and errors from either the PRINCE and TAFKAP decoders.

**Figure 4:**
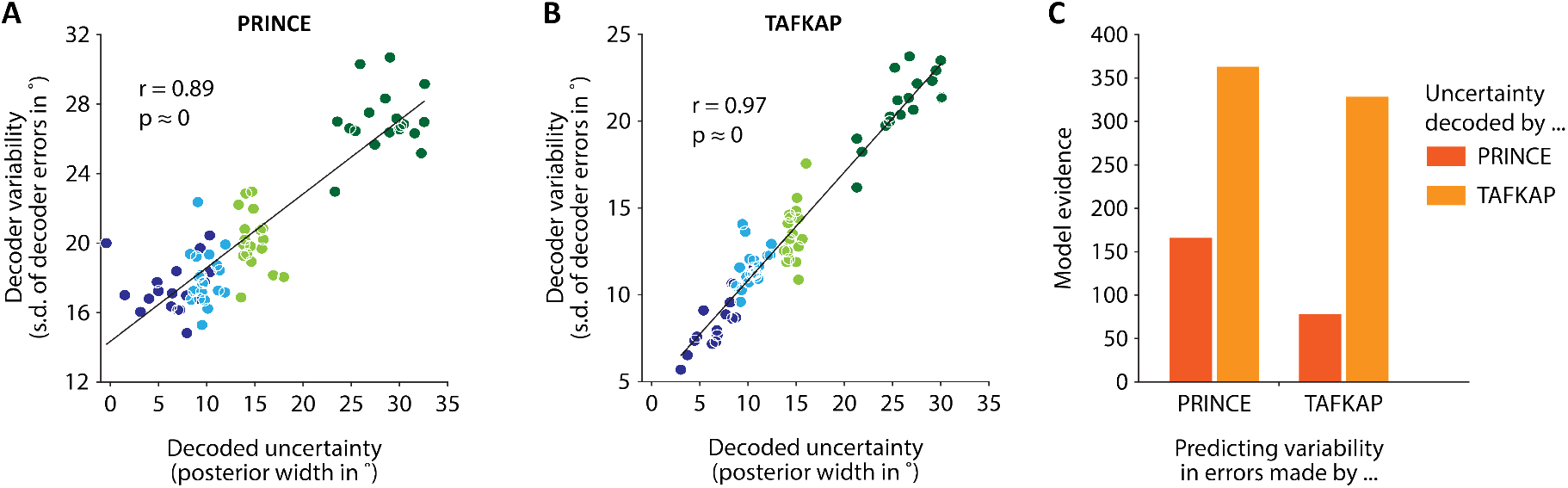
Uncertainty estimates from the new TAFKAP decoder better reflect the precision of information in cortical data. Panels (**A**) and (**B**) show the correlation between trial-by-trial uncertainty estimates and the variability of the decoded stimulus estimates, for PRINCE and TAFKAP respectively. Dots indicate the mean decoded uncertainty and variability in decoded estimates for each of four uncertainty bins per participant (bins shown in different colors). Lines show the best linear fit to the data. Panel (**C**) shows the results of a bin-free analysis, quantifying the degree to which decoded uncertainty predicts the error in decoded estimates of either algorithm (on the same trials). Bars show the evidence (negative BIC) for a link between decoded uncertainty and error variability.

**Figure 5:**
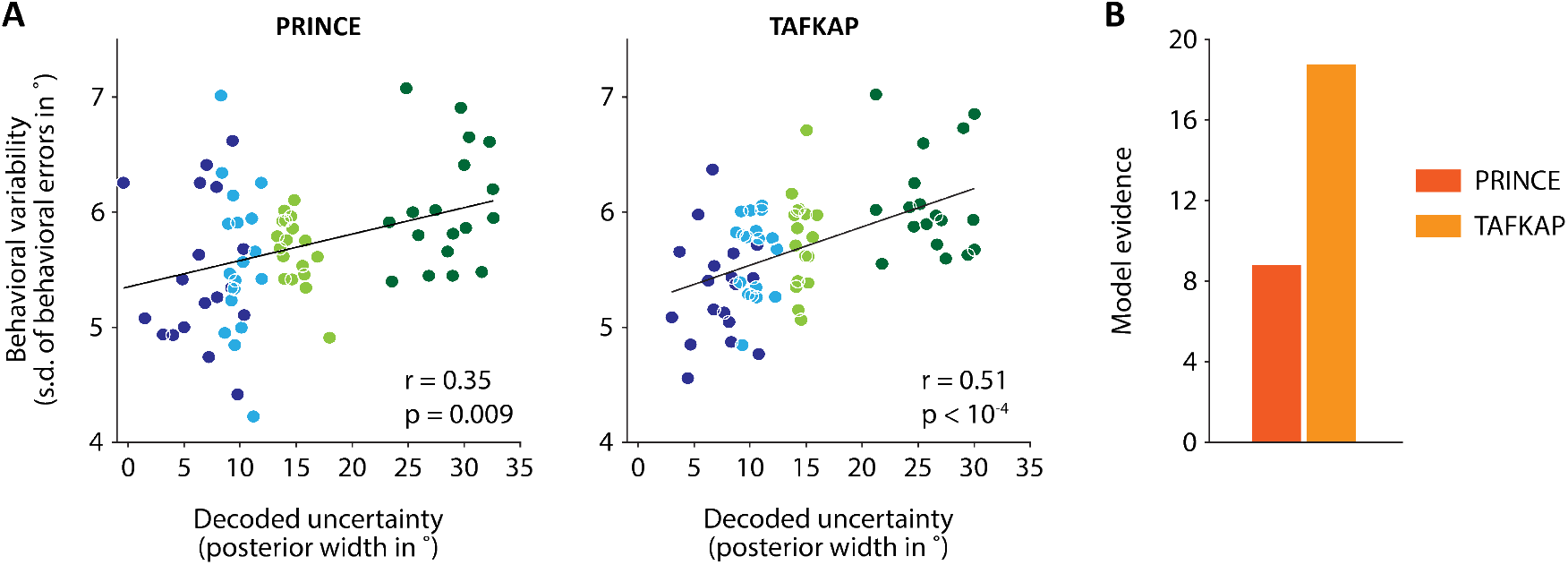
Uncertainty estimates from TAFKAP better predict human behavior. (**A**) Behavioral variability plotted against decoded uncertainty, in four uncertainty bins per participant, for both PRINCE and TAFKAP. One dot is plotted for each bin and participant, with bins indicated by different colors. Lines show the best linear regression fit. A numerically stronger correlation is apparent for the new decoder. (**B**) Model evidence (negative BIC) for a trial-by-trial link between decoded uncertainty and the width of behavioral error distributions, as compared to a null model in which no such link exists. The stronger evidence for the new decoder (difference in BIC of 9.9) offers strong statistical support for the notion that the new decoder better predicts behavior.

### Code availability

Custom Matlab code for the PRINCE and TAFKAP probabilistic decoding methods is publicly available, and can be found at https://github.com/jeheelab/TAFKAP

## Results

This paper introduces two measures to improve decoding of probabilistic stimulus representations from cortical activity patterns: (1) resampling the training data and (2) shrinkage-based estimation of the spatial noise covariance matrix. We will first discuss the rationale behind these two measures, and demonstrate that they improve decoding performance for simulated data. Next, we will apply the algorithm to fMRI data from human participants to find that decoding performance of both the orientation stimulus and its associated uncertainty in cortex is greatly improved.

### Theory & simulations

The probabilistic decoding approach upon which we build here is discussed in detail elsewhere^5,8^ as well as in the Methods section. Its basic principles are as follows. We start by specifying a *generative model* that describes how a visual stimulus leads to a measurable cortical response. Knowing this relationship will later enable us to reason backwards from an observed pattern of cortical activity to the stimuli that may have caused it (**Fig. 1A**). For convenience, we will assume in what follows that cortical activity is measured with fMRI, and that the stimulus feature of interest is its orientation. However, the same approach can also be applied to other types of neural data and stimulus features.

In the generative model specified here, the fMRI BOLD response to a stimulus consists of two components: a tuning function *f*(*s*) and random variability. The tuning function describes the voxel’s mean response to repeated presentations of stimulus *s*. Random variability (or noise) corresponds to the across-trial fluctuations in this response. The noise in each voxel’s response is assumed to be Normally distributed, with voxel-by-voxel covariance matrix **Ω**. Thus, the probability of observing a cortical activity pattern **b** given stimulus orientation *s* is given by a Normal probability model:

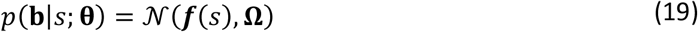

where ***f***(*s*) = [*f*_*i*_(*s*)] is a vector containing each voxel’s tuning curve value for stimulus *s*, 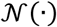 denotes the (multivariate) Normal probability density function, and **θ** is the set of all free parameters in the model (i.e. voxel noise covariances and tuning curves), which are estimated from training data (**Fig. 1C**).

The goal of the decoding algorithm is to recover the posterior probability distribution *p*(*s*|**b**), which describes how likely it is that the stimulus had orientation *s* given the observed cortical activity pattern **b**. This posterior can be computed by applying Bayes’ rule:

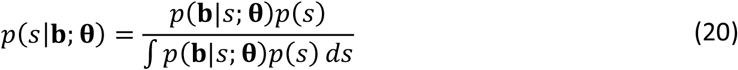

where *p*(*s*) describes, for each stimulus, the a-priori probability that it was presented. Thus, the posterior is computed by *inverting* the generative model, reasoning back from an observed activity pattern to the stimuli that could have evoked it (**Fig. 1A**).

The posterior depends on the parameters of the generative model, **θ**. The best possible decoder uses the true values of these parameters (e.g., it accurately models the voxel tuning functions, and the amount of (correlated) noise in the data, with no measurement error) to calculate the posterior from a pattern of cortical activity. The variance or width of this posterior can be taken as a measure of the uncertainty contained in cortical activity pattern **b** about the stimulus *s*. We will refer to this as *intrinsic uncertainty*: the degree of uncertainty inherent to any pattern of noisy activity, even when using the true value of model parameters. A main goal for probabilistic decoding of neuroimaging data is to estimate this intrinsic uncertainty as accurately as possible.

In practice, however, the true values of the generative model parameters are unknown. Instead, they are estimated from limited and noisy training data, which makes the parameter values *themselves* uncertain. As a consequence, these imprecise parameter values introduce additional uncertainty in the interpretation of any pattern of cortical activity, which we refer to as *model uncertainty*. This paper addresses this model uncertainty in two ways. First, we average posteriors across many plausible model settings, and show that such averaging improves decoding accuracy. Second, we introduce improved methods for estimating the voxel noise covariance matrix, which we show is more flexible than a previous implementation and also results in better decoding performance.

#### Bagging to account for model uncertainty

How to account for uncertainty that arises due to imprecise model parameter estimates? Previous decoding algorithms^5,6^ used a single “best guess” estimate of the generative model parameters – for instance, the value of **θ** that maximizes the probability of the training data. While this is likely the most accurate single (point) estimate of the model parameters, the value alone does not reflect the full scope of values that are plausible under the training data. To better account for model uncertainty (and thereby total uncertainty), we here proposed the following solution: randomly resample the training data for many iterations, find the best fitting model parameter values for each instance, and then average the obtained posterior distribution across iterations (**Fig. 1D**). Specifically, by repeatedly sampling observations (i.e. pairs of activity patterns and stimulus labels) at random and with replacement from the training data, and computing best-guess estimates of the model parameters from each resampled data set, a large set of plausible tuning curves and noise covariances can be constructed. Each of these parameter samples **θ**^∗^ is then used to compute a stimulus posterior for the same activity pattern in the test set (**b**_test_). Finally, these distributions are averaged into a single decoded posterior – a step akin to mathematical marginalization. The so-obtained posterior no longer depends on a particular value of **θ**, but only on the training data which allowed us to obtain samples of **θ**. This type of procedure is also known as bootstrap-aggregating or “ bagging”^9^. An important advantage of bagging is that it does not require many assumptions and automatically follows the empirical distribution of the data.

To validate this bagging procedure and its effect on decoding performance, we used synthetic data for which the true (encoded) stimulus posteriors were known by construction. fMRI data sets were simulated for 18 model participants, with realistic noise covariance structures modeled after those for actual participants, and divided into a train and test set. We then trained and tested two decoders on this data: the *PRINCE* decoder, which uses best guess-estimates of the generative model parameters, and *PRINCE+bagging*, which implements the bagging-based procedures described above. For each activity pattern in the test set, we thus obtained three distributions: the true posterior (using the parameters with which the data were generated), and the posteriors from the two decoding methods, with and without bagging. We evaluated the accuracy of the decoded posteriors by computing their Kullback-Leibler divergence to the true posterior (low KL-divergence indicates high similarity). Interestingly, we found that PRINCE+bagging yielded significantly better results than PRINCE (**Fig. 2A**; paired *t*-test on the difference in KL-divergence to the ground truth posterior: *t*(17) = 7.5, *p* < 10^−6^). These results demonstrate that bagging can serve as a cheap and effective way to more accurately estimate the posterior distribution encoded in cortical activity.

**Figure 2:**
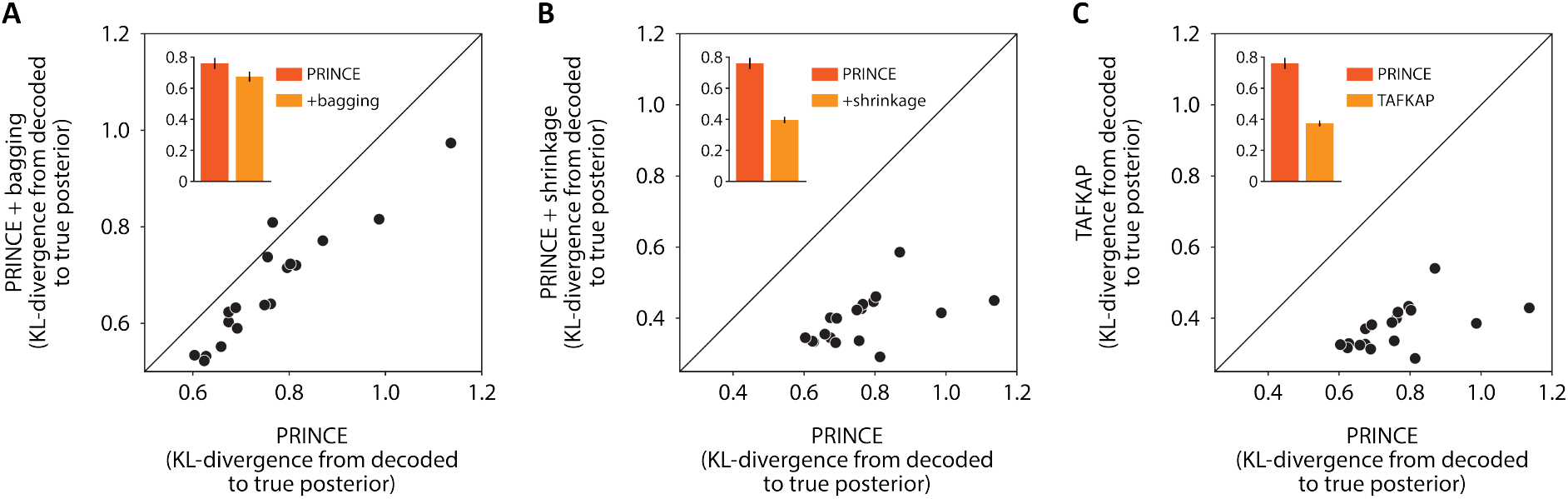
Simulations indicate that probabilistic decoding can be improved by incorporating the two proposed measures, sampling (or *bagging*) and shrinkage. (**A**) Bagging leads to more accurate decoded posteriors (lower KL-divergence to the true posterior) compared to the PRINCE decoder, which uses best-guess estimates of the generative model parameters. (**B**) A flexible shrinkage-based estimation procedure for the noise covariance structure also improves the accuracy of the decoded posterior. This method blends the (complex, but error-prone) sample covariance matrix with a simpler target structure. This is shown to perform better than the approach used in the PRINCE decoder, which used the simpler covariance structure alone (and thus effectively used a fixed shrinkage towards this target of 100%). (**C**) The two measures combined result in the new decoding algorithm TAFKAP, which outperforms PRINCE, PRINCE+bagging and PRINCE+shrinkage. In all panels, dots show the mean KL-divergence between decoded and true posteriors for 18 simulated subjects (lower KL-divergence is better). Standard errors for individual simulated subjects were too small to display. Bars and error bars in insets show across-subject averages +/− 1 SEM.

#### Shrinkage-based covariance estimation

The noise covariance structure across voxels can be especially challenging to estimate from limited training data. The most straightforward method is to simply compute the sample covariance of the residual voxel responses around the fitted tuning curves:

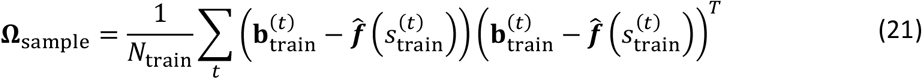

where 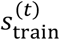 and 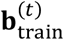 are the stimulus and activity pattern, respectively, for trial *t* in the training set. This sample covariance is also the maximum likelihood estimator of **Ω**. However, when there are more voxels in the training data than there are observations (trials), the sample covariance matrix is not invertible. When this happens, the posterior is undefined, and so decoding is impossible. Moreover, even when trials outnumber voxels, the ratio of data to voxels is small enough in typical fMRI data sets to make the sample covariance a very unstable estimator, with large imprecision and estimation errors. This is a well-known problem in statistics, and the typical solution, called *shrinkage estimation*^10^, is to blend the sample covariance with an estimator that is more stable by virtue of having fewer free parameters (**Fig. 1E**):

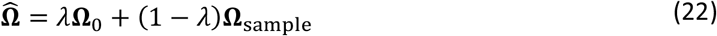

where **Ω**_0_ is the simpler target to which the sample covariance is “shrunk”, and 0 ≤ *λ* ≤ 1 is the degree of regularization applied. Previously, we implemented an extreme version of this, which used only the simpler structure **Ω**_0_ to capture voxel noise correlations (effectively setting *λ* to 1). The covariance structure we use as a shrinkage target is conceptually given by:

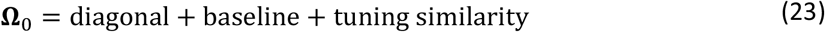

That is, we assume that voxel noise is a combination of independent noise (modeled by a diagonal of independent noise variances), noise that is correlated between all voxels (modeled by a baseline correlation), and noise that is shared specifically between voxels with similar tuning curves (see Methods for more detail). While this structure captures the most important aspects of the voxel noise covariance for decoding^8^, some residual variance not modeled by the structure may nonetheless be relevant to extracting stimulus information from cortex. To be sensitive to such additional variation, the new TAFKAP algorithm lets the sample covariance also contribute to 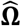. The weight of the sample covariance is determined by finding the optimal value of *λ* using a cross-validation procedure within the training data (see Methods). This approach thus aims to find the optimal middle ground between stability and flexibility.

We tested the degree to which the modified covariance structure improved decoding performance using the same synthetic data that was used to assess the effect of bagging. For each simulated activity pattern, we again computed the ground-truth posterior (using the parameters by which the data were generated), the posterior decoded using the PRINCE method (which uses a fixed shrinkage strength of *λ* = 1), and the posterior decoded by PRINCE+shrinkage, where the degree of covariance shrinkage is tailored to the data. With PRINCE+shrinkage, the mean KL-divergence to the true posterior was found to be significantly smaller than with the original PRINCE method (**Fig. 2B**; paired *t*-test on the difference in KL-divergence to the true posterior: *t*(17) = 13.4, *p* < 10^−9^). This provides a proof-of-principle that blending the sample covariance with the previously developed low-parameter structure can improve the accuracy of decoded stimulus posteriors.

#### TAFKAP: combining bagging and shrinkage

Finally, we used the synthetic fMRI data to evaluate the effect of augmenting the PRINCE decoding method with both bagging and (flexible) covariance shrinkage, to verify that these two measures would work well together. We call the resulting decoding method *TAFKAP;* **T**he **A**lgorithm **F**ormerly **K**nown **A**s **P**rince. Posteriors decoded using this method were even more faithful to the true (simulated) posteriors, than when PRINCE was augmented with bagging or shrinkage alone (TAFKAP vs. PRINCE: *t*(17) = 14.4, *p* < 10^−10^; TAFKAP vs. PRINCE+bagging: *t*(17) = 12.7, *p* < 10^−9^; TAFKAP vs. PRINCE+shrinkage: *t*(17) = 7.75, *p* < 10^−6^).

### Human fMRI data

Having demonstrated the validity of the TAFKAP algorithm on synthetic data, we now turn to fMRI data collected from human observers to determine whether TAFKAP also decodes more accurate posteriors from cortical activity in practice. BOLD activation patterns were measured in early visual cortex (areas V1, V2 and V3) while participants viewed randomly oriented grating stimuli. A few seconds after each stimulus was presented, the participant reported its orientation by rotating a bar presented at fixation. The activity pattern evoked by each stimulus was run through both the PRINCE and TAFKAP versions of the probabilistic decoding algorithm. Thus, we obtained two decoded posterior distributions for each trial in the experiment: one for each decoder. We then asked the question: which of these distributions better reflects the information in 1) the recorded BOLD activity patterns, and 2) the underlying neural responses?

When analyzing real fMRI data, there is no gold-standard ‘true posterior’ to which decoded posteriors can be directly compared, as is possible for simulated data. We therefore relied on two different metrics, which indirectly quantify the accuracy of the decoded distributions: first, how well the decoded stimulus estimates (derived from these distributions) correspond to the actual presented stimulus orientations, and second, the degree to which the width of the decoded distributions predicts the errors in these orientation estimates. In addition, we also examined how well decoded uncertainty predicted the variability in participant’s behavior. This final analysis addresses the extent to which the decoded distributions characterize the trial-by-trial quality of information encoded in neural stimulus representations, as explained in more detail below.

#### Orientation decoding performance

While orientation decoding performance does not capture the full gamut of information in the activity pattern, it is still an important benchmark, as a decoder can only improve on this task by making better use of the available information. For both decoding algorithms, the stimulus estimate for a given trial is obtained by taking the (circular) mean of the decoded posterior on that trial (i.e. the center of probability mass; **Fig. 1B**). We evaluate orientation decoding performance in two ways: by calculating the mean absolute error with respect to the true stimulus orientation, and by computing the circular correlation between the decoded and actual stimulus orientations. We find that TAFKAP performs better than PRINCE on both of these metrics (**Fig. 3A&C**). Compared to the PRINCE algorithm, the mean error in the decoded estimates of TAFKAP decreased by approximately 36%, from 17.4° to 11.1°, and this improvement was highly significant for each observer in the experiment (Wilcoxon signed rank tests, least significant participant: *Z* = 5.3, *p* = 1.2 × 10^−7^), as well as across observers (*Z* = 3.7, *p* = 2.0 × 10^−4^). Correlations between presented and decoded orientations increased from 0.73 to 0.88, and this too was a statistically significant improvement, both across participants and for each individual (least significant *p*-value from permutation tests for individual observers: *p* = 5.9 × 10^−3^; Wilcoxon signed rank test across observers: *Z* = 3.7, *p* = 2.0 × 10^−4^). These results demonstrate that TAFKAP extracts substantially more of the stimulus information available in cortical activity than PRINCE. Interestingly, when evaluated across subjects, the best decoding performance attained by TAFKAP even rivaled that participant’s behavioral performance: TAFKAP attained a mean absolute decoder error of 5.7°, while that same participant’s behavioral accuracy was 6.8° (mean absolute behavioral error before bias correction).

**Fig. 3:**
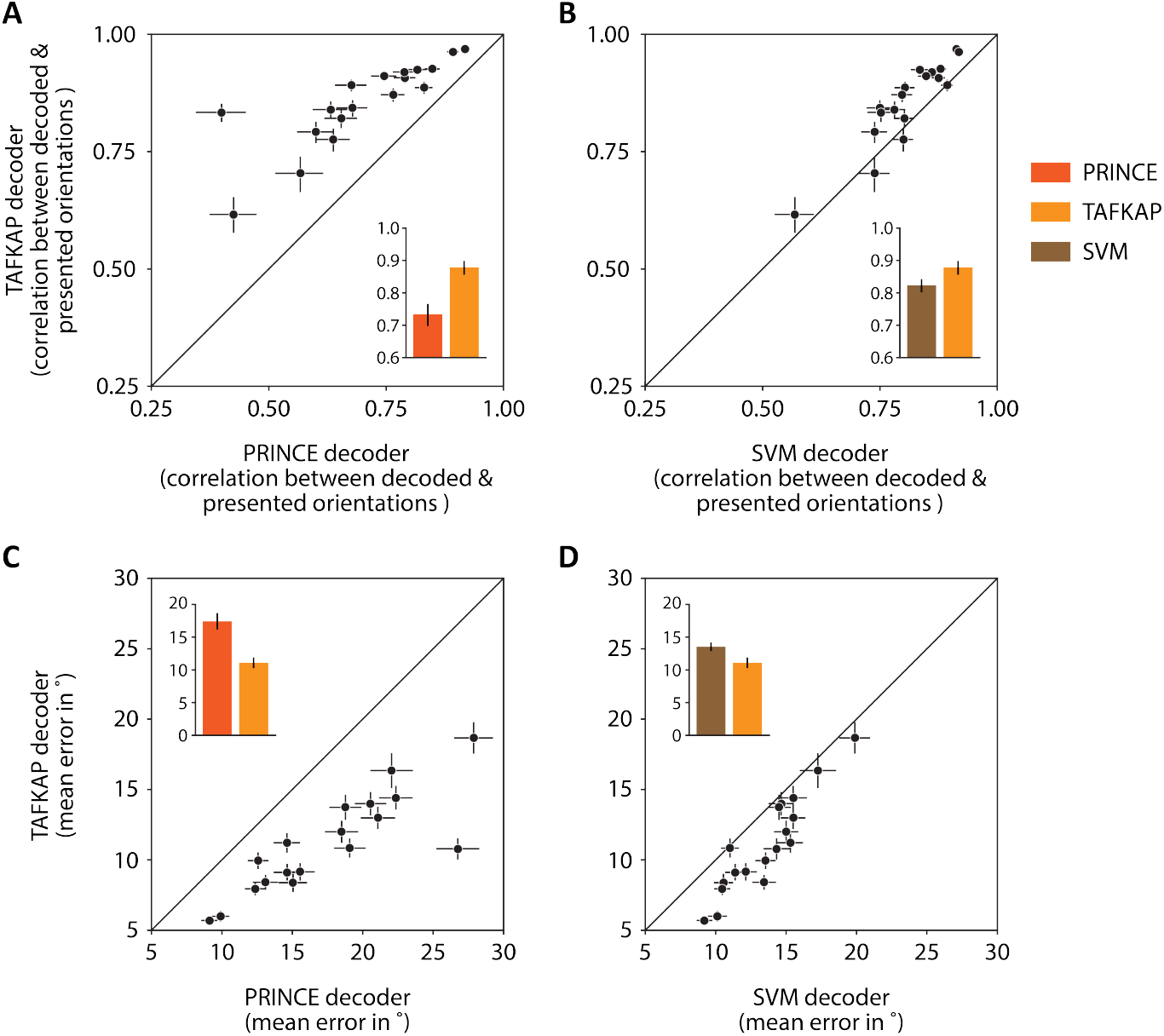
Improved decoding of stimulus orientations from activity in human visual cortex. (**A-D**) TAFKAP produces better estimates of the presented stimulus compared to PRINCE, and compared to an SVM decoder. This is indicated by a stronger correlation (**A-B**) and smaller errors (**C-D**) between decoded and presented orientations. Dots correspond to values for individual observers, while insets show averages across participants. Error bars indicate +/− 1 SEM.

Decoding a ‘best guess’-estimate of the presented stimulus can also be achieved with other algorithms, and is widely used in the field. Our decoding approach differs from such previous approaches in that it provides an estimate of the trial-by-trial quality of information contained in the cortical population response, rather than a point estimate of the stimulus – a feature important for neuroimaging studies of Bayesian decision-making^5,6^. Nevertheless, it is interesting to see how our method compares to these more conventional decoders. One of the most popular classes of algorithms to decode neuroimaging data is provided by Support Vector Machines. We therefore compared orientation decoding performance from PRINCE and TAFKAP to that of an SVM trained and tested on our fMRI data. Do we lose out on ‘best guess-accuracy’ by decoding probability distributions? For PRINCE, the answer to this question is affirmative, as the SVM decoder achieved a mean error of 13.5° and a correlation of 0.82, both of which were significantly better than PRINCE (Wilcoxon signed rank tests, *Z* = 3.1, *p* = 0.002 and *Z* = 3.5, *p* = 4.6 × 10^−4^, respectively). TAFKAP, on the other hand, produced more accurate orientation estimates than the SVM, despite not being optimized for this purpose (**Fig. 3B&D**; Wilcoxon signed rank tests, *Z* = 3.7, *p* = 2.1 × 10^−3^ and *Z* = 3.3, *p* = 8.6 × 10^−4^, for the mean error and correlation between decoded and presented orientations, respectively). This is noteworthy, as it suggests that researchers can choose to use probabilistic decoding without making concessions on best-guess decoding accuracy.

#### Uncertainty decoding performance

This improved estimation of the presented stimulus indicates that the mean of the decoded distribution is more accurate for the new decoder. But does the uncertainty in the decoded distributions (i.e. their width) also better reflect the fidelity with which stimuli are represented in cortical activity patterns? In our simulation analyses, we answered this question by comparing the decoded probability distributions to the true distributions that were known to be encoded in the simulated data. For actual neuroimaging data, this approach cannot be used, as the true distributions are unknown (otherwise, there would be no need for decoding algorithms). We must therefore rely on less direct methods to verify that decoded uncertainty reflects the information in cortical activity.

What metric can be used to verify that decoded uncertainty reflects the actual uncertainty in the data? If an activity pattern truly has high uncertainty, then an estimate of the stimulus presented on that trial should (on average) be less precise. That is, the orientation estimates decoded from such patterns will follow a wider distribution around the true stimulus orientation. On the other hand, activity patterns with low uncertainty should lead to more accurate stimulus estimates. Thus, if decoded uncertainty reflects actual uncertainty, then decoded uncertainty should reliably predict the accuracy of decoded stimulus estimates.

To examine this relationship, trials were divided into four bins (separately for each observer) of increasing decoded uncertainty. Across trials in each bin, we calculated the mean decoded uncertainty, and the circular standard deviation of the errors in the decoder’s orientation estimates. Across bins and participants, we then computed the partial correlation between decoded uncertainty and error variability. As expected based on our previous work^5^, this correlation was clearly visible and highly significant for both decoders (**Fig. 4A-B**).

While this indicates that both PRINCE and TAFKAP can reliably characterize the degree of uncertainty in cortical activity patterns, which of the two algorithms performs better? Although for PRINCE the correlation between decoded uncertainty and the variability of orientation errors was numerically higher, note that these are correlations between separate pairs of variables, as both the decoded uncertainty values and estimation errors are different between decoders. Thus, it could be that PRINCE and TAFKAP both decode uncertainty equally well, while the orientation estimates are better (more predictable) for TAFKAP, resulting in a numerically larger correlation coefficient. In addition, while the binning procedure is helpful in order to visualize the predicted relationship, ideally one would like to perform the analysis without this intermediate step. To address these two issues, we devised an additional analysis that does not require binning, and which allowed uncertainty from one decoder to be used to predict the errors of the other. This analysis was carried out on the raw, trial-by-trial uncertainty values and decoder errors, both within and between decoders. Rather than binning data in order to compute an across-trial measure of error variability, we explicitly modeled the distribution of the decoder errors. Specifically, trial-by-trial data were fit with a model in which decoder errors followed a Normal distribution, with a standard deviation that increased with decoded uncertainty.

Does this model-based analysis bear out the earlier results, suggesting that TAFKAP better extracts the uncertainty in fMRI activity patterns? If so, we should observe stronger evidence for the models that use TAFKAP’s uncertainty to predict decoder errors. Interestingly, we found that compared to PRINCE, TAFKAP was indeed much better able to predict its own errors, as well as those of the PRINCE algorithm (Fig**. 4C**). In both cases, model evidence (negative BIC – see Methods) was over 200 points higher for TAFKAP, indicating very strong statistical support^23^ for this new approach over PRINCE. Since TAFKAP also performs better when its decoded uncertainty values are used to predict PRINCE’s decoding errors, this effect cannot be attributed simply to better orientation estimates, and must really be due to more accurate uncertainty estimates. This suggests that TAFKAP estimates the uncertainty in cortical activity substantially better than PRINCE.

As foreshadowed by our simulation results, the empirical findings so far show that the new updates to our decoding algorithm lead to a substantially better characterization of the information contained in cortical activity patterns. But do the decoded distributions also better characterize *neural* information? This is not a given. Neuroimaging measurements are subject to additional sources of noise and ambiguity, on top of the underlying neural response. Decoded uncertainty might reflect these non-neural sources more than it tracks the uncertainty in the neural response, and only the latter is of interest to neuroscientists.

How can neural and non-neural sources of uncertainty be disentangled? Note that neural uncertainty impacts behavior, whereas uncertainty that derives from the neuroimaging method cannot possibly influence behavioral stimulus estimates. Specifically, a high-quality, unambiguous neural representation of a stimulus allows the observer, on average, to more accurately report its orientation. In contrast, a noisy, highly uncertain neural stimulus representation limits the observer’s performance, and will (on average) lead to less precise behavior. Thus, if decoded uncertainty reflects neural uncertainty, it should predict the precision of behavioral stimulus reports. To quantify the correlation between decoded uncertainty and behavioral precision, we therefore employed the same analysis methods that we used to correlate decoded uncertainty with the errors in the decoded stimulus orientation. Instead of decoded orientation estimates, we now looked at the relationship between decoded uncertainty and the errors in behavioral orientation estimates. A stronger correlation is consistent with a closer link between decoded and neural uncertainty.

Is TAFKAP more sensitive to neural uncertainty than PRINCE? Interestingly, our findings indicate that TAFKAP does produce better estimates of the degree of uncertainty contained in neural population activity. For TAFKAP, the correlation between decoded uncertainty and behavioral error variability (computed across four uncertainty bins per participant) was numerically higher than for PRINCE (increasing from *r* = 0.35, *p* = 0.009 to *r* = 0.51, *p* < 10^−4^; **Fig. 5A**). This result was statistically validated by a bin-free model comparison: the evidence (negative BIC) for a link between decoded uncertainty and behavioral precision was 9.9 points higher for TAFKAP than for PRINCE (**Fig. 5C**), indicating strong statistical support^23^. This finding is especially exciting, as it suggests that a few easily implemented modifications have rendered the decoding method substantially more sensitive to the trial-by-trial fidelity with which sensory information is represented in cortex.

## Discussion

We have presented two modifications to our previously-developed probabilistic decoding method, which markedly improve the method’s ability to characterize the information contained in cortical activity patterns. Using bagging to average over many plausible settings of the generative model parameters, combined with shrinkage-based estimation of the voxel noise covariance, the new algorithm achieved better decoding performance than the existing approach. Estimates of the presented stimulus from the new TAFKAP method were substantially more accurate than those of its predecessor PRINCE, and also better than an SVM-based decoder. Moreover, the uncertainty estimates also correlated more strongly with decoder and behavioral errors than before, suggesting that the distributions decoded by TAFKAP more faithfully reflect the information in cortical stimulus representations.

The advent of multivariate “decoding” methods in neuroimaging analysis, nearly two decades ago^24–26^, opened a new window for researchers to examine the information contained in cortical activity patterns. These methods can, for instance, provide an estimate of the sensory stimulus that most likely evoked a certain cortical response. But neuroscientists have long been aware that, due to various sources of random variability or noise, cortical activity is typically consistent with a range of possible interpretations, rather than just a single value^27^. Probabilistic decoding methods allow brain researchers to tap into this full gamut of possibilities, and thus reveal the precision or uncertainty with which a particular stimulus is represented. This has several important benefits over the single-value decoding approach.

First, decoding probabilities allows us to test probabilistic (or Bayesian) theories of neural representation and computation. Such theories posit that probabilities are the representational currency used by the brain to encode sensory information^4,28^. Furthermore, they predict that observers should adjust their perceptual decisions based on the uncertainty associated with different sources of sensory evidence. Probabilistic decoding methods enable one to validate these predictions against the behavior and cortical representations of human observers. Notably, using our PRINCE algorithm, we previously showed that the degree of uncertainty in cortical stimulus representations predicts, on a trial-by-trial basis, how much weight observers give to this information in their decisions. Interestingly, some of our findings were recently replicated in macaque visual cortex, similarly using a probabilistic decoding algorithm^7^. Thus, probabilistic decoding methods have strengthened the case for Bayesian theories of neural coding, in ways that would not have been possible otherwise.

Second, decoding probabilities can also be advantageous when probabilistic neural representations are not the primary focus. A decoded probability distribution not only gives a ‘best-guess’ estimate of what is being decoded (e.g. stimulus orientation), but also provides a measure of how confident one ought to be in this estimate. Thus, if a particular analysis requires very precise estimates, probabilistic decoding can help eliminate trials that do not meet the required threshold of reliability. One interesting application of this principle may be in brain-computer interfaces (BCIs), which control robots or other electronic devices through neural activity. It may be desirable for a BCI to only initiate an action if the recorded neural signals are sufficiently unambiguous. Probabilistic decoding provides a natural way to quantify this ambiguity, and incorporating the approach might help create more reliable BCIs.

The alterations in probabilistic decoding methods that we have presented here are fairly straightforward to implement, while yielding substantial improvements. Moreover, our results suggest that choosing probabilistic decoding over more established methods, such as SVMs, does not come at the expense of decoding accuracy. TAFKAP, in fact, can produce more accurate orientation decoding performance than an SVM, and in some cases even approached the participant’s behavioral accuracy. Moreover, the method can easily be applied to decode other continuous variables from visual cortex, and it can be adapted to decode discrete variables, target different brain areas, or decode data from other imaging modalities. TAFKAP, then, is not meant to only be a specialized tool for a specific purpose. In many cases, it can be used in place of a conventional decoder with little effort, while getting probabilities and uncertainty as an additional dimension in the analysis.

In summary, probabilistic decoding is a novel tool in the arsenal of brain researchers that has the potential to open up exciting, new avenues for neuroimaging studies. Here, we have presented TAFKAP: an improved probabilistic decoding method that builds on previous work^5^. Our findings indicate that the probability distributions decoded by TAFKAP more closely reflect stimulus information in fMRI activity patterns, including the degree of uncertainty contained in neural population activity. “Best-guess” stimulus decoding was also more accurate, and competitive with an SVM-based decoder. We hope that the current results will inspire more researchers to add probabilistic decoding to their belt of analysis tools.

